# Neural phoneme discrimination in variable speech in newborns – associations with dyslexia risk and later language skills

**DOI:** 10.1101/2023.01.10.522962

**Authors:** P. Virtala, T. Kujala, E. Partanen, J. A. Hämäläinen, I. Winkler

**Affiliations:** Cognitive Brain Research Unit, Centre of Excellence in Music, Mind, Body and Brain, Department of Psychology and Logopedics, Faculty of Medicine, University of Helsinki, Finland; Jyväskylä Centre for Interdisciplinary Brain Research, Department of Psychology, University of Jyväskylä, Finland; Institute of Cognitive Neuroscience and Psychology, Research Centre for Natural Sciences, Budapest, Hungary

**Author notes:** shared first authorship (P. Virtala, T. Kujala). **Corresponding author:** Dr. Teija Kujala, Cognitive Brain Research Unit, P.O. Box 21, 00014 University of Helsinki, Finland.

**Keywords:** Infants, dyslexia, mismatch responses (MMRs), phoneme processing, language learning

## Abstract

A crucial skill in infant language acquisition is learning of the native language phonemes. This requires the ability to group complex sounds into distinct auditory categories based on their shared features. Problems in phonetic learning have been suggested to underlie language learning difficulties in dyslexia, a developmental reading-skill deficit. We investigated auditory abilities important for language acquisition in newborns with or without a familial risk for dyslexia with electrophysiological mismatch responses (MMRs). We presented vowel changes in a sequence of acoustically varying vowels, requiring grouping of the stimuli to two phoneme categories. The vowel changes elicited an MMR which was significantly diminished in infants whose parents had the most severe dyslexia in our sample. Phoneme-MMR amplitude and its hemispheric lateralization were associated with language test outcomes assessed at 28 months, an age at which it becomes possible to behaviourally test children and several standardized tests are available. In addition, statistically significant MMRs to violations of a complex sound-order rule were only found in infants without dyslexia risk, but these results are very preliminary due to small sample size. The results demonstrate the relevance of the newborn infants’ readiness for phonetic learning for their emerging language skills. Phoneme extraction difficulties in infants at familial risk may contribute to the phonological deficits observed in dyslexia.

**Research highlights:**

- We recorded mismatch responses (MMRs) to vowel changes in a variable speech stream
- Newborns extracted relevant phonetic information from the stream, evidenced by MMRs
- High dyslexia risk infants had diminished MMRs to vowel changes
- MMR amplitudes and hemispheric lateralization correlated with later language skills
- Poor phoneme extraction may compromise phonological and language development

## 1. Introduction

Native language acquisition via auditory learning begins at the earliest stages of infancy, and involves, for example, the adoption of native language phoneme categories (Kuhl, 2010; Serniclaes, 2018). Atypical development due to heritable disorders, such as developmental dyslexia (DD; a disability in age-appropriate reading-skill acquisition, Peterson and Pennington, 2015), may lead to difficulties in subsequent language and literacy acquisition (Eden et al., 2016; Giraud and Ramus, 2013; Vellutino et al., 2004). A range of perceptual-cognitive deficits have been associated with DD (e.g., Peterson and Pennington, 2015), including phonological (e.g., Eden et al., 2016; Snowling and Melby-Lervåg) and temporal-processing deficits (potentially underlying the phonological deficit; e.g., Kalashnikova et al., 2021). According to the leading theory, inaccurate phoneme representations or poor access to them, i.e., phonological deficit, is the main cause of DD (Eden et al., 2016; Giraud and Ramus, 2013; Vellutino et al., 2004; see Snowling and Melby-Lervåg, 2016, for a meta-analysis). Here, our main aim was to assess newborn auditory abilities potentially important for language acquisition and the development of DD, namely, native language phoneme discrimination as an index of phonetic learning, and its association with familial DD risk and subsequent language skills. Better understanding of infant neurocognitive impairments associated with language and reading disorders could help in revealing their developmental trajectories, and, should these associations reflect causal connections, in designing effective preventive interventions.

So far, event-related potential (ERP) studies on infants have shown that well-functioning auditory abilities promote efficient language development (Ramus, 2002), as they are associated with future language and literacy measures and can be compromised by familial DD risk (for meta-analyses: Oh et al., 2019; Volkmer and Schulte-Koerne, 2018). For example, typically developing newborn infants can detect consonant changes in syllables even when they are uttered by several different speakers (Dehaene-Lambertz and Pena, 2001) and order changes in paired sounds, when these pairs vary over several frequency levels (Carral et al., 2005; see also Háden et al., 2015 and Stefanics et al., 2009). These abilities to discriminate sound features in an acoustically varying context suggest that neonatal auditory skills extend to categorizing sounds, even based on abstract rules on interstimulus relationships. This is thought to be a core ability for language acquisition (Kuhl, 2010), for example for learning phonotactics and morphosyntax, as well as for forming accurate phoneme representations. However, auditory processing deficits in familial DD risk, for example in phoneme discrimination, have often been studied with repetitive, acoustically non-varying stimulation streams, which do not necessarily require the ability to categorize sounds based on shared features (Oh et al., 2019; Volkmer and Schulte-Koerne, 2018). In our recent adult studies, we attempted to tackle this issue by presenting phoneme changes and violations of an abstract rule in an acoustically varying context (Virtala et al., 2021, 2020). They both elicited diminished neural responses in DD (Virtala et al., 2021, 2020), and the deficient rule extraction tended to be associated with poor reading performance assessed in this study (Virtala et al., 2021). Crucially, phoneme changes in an acoustically non-varying, repetitive context did not demonstrate group differences (Virtala et al., 2020). Finding diminished responses in the acoustically variable condition and not in the repetitive condition suggests that phoneme categorization rather than the discrimination of the acoustic differences between the phonemes was deficient in DD. This supports the observation that basic acoustic discrimination deficits are not always present in DD (review: Hämäläinen et al., 2013). Furthermore, the findings suggest that ERPs elicited by complex auditory streams requiring categorizing the stimuli based on shared features and previous experience (native language learning) may be particularly sensitive measures of DD and its familial risk. However, to our knowledge, no previous study has tested the ability to extract phonetically relevant information from natural-like acoustic variation in infants at DD-risk, or the associations of these abilities to subsequent language skills.

In the current study, we examined newborn native language phoneme discrimination in an acoustically variable context as a potential index of crucial language learning abilities. We employed the ERP paradigm we previously used in adults with or without DD (Virtala et al., 2021, 2020), and measured neonatal mismatch responses (MMRs; Kushnerenko et al., 2013), an ERP component elicited even in sleeping newborns and regarded as the neonatal counterpart of the adult mismatch negativity (MMN; Näätänen et al., 2019). MMN is a negatively-displaced ERP at fronto-central channels to acoustic deviation and considered an index of stimulus discrimination accuracy (Kujala and Näätänen, 2010). It can reflect the processing of phonemes (Näätänen et al., 1997) as well as abstract rule extraction (Paavilainen, 2013). The MMN amplitude correlates with behavioral discrimination accuracy (Tiitinen et al., 1994; Virtala et al., 2018). It is larger for native than for foreign phoneme contrasts and thus reflects phonetic learning (Winkler et al., 1999). In children and adults, MMNs to speech sounds are usually more left-ward lateralized relative to MMNs to nonspeech sounds (Kuuluvainen et al., 2016; Sorokin et al., 2010). MMRs in newborns are often of positive polarity (Kushnerenko et al., 2013), but also negative polarities have been reported in infancy (e.g., Partanen et al., 2013; Thiede et al., 2019), particularly to acoustically salient deviants (e.g., Cheng and Lee, 2018). For example, in our previous study, vowel deviants only elicited a P-MMR, while acoustically more salient duration and frequency deviants also elicited negativities in healthy control infants (partly the same infants as in the present study; Thiede et al., 2019). Indeed, these responses may also co-exist, with negative MMRs typically seen at an earlier latency than the positive MMRs (e.g., Fellman et al. 2004; Partanen et al., 2013; Thiede et al., 2019).

It is noteworthy that in previous infant-MMR studies on the effects of familial DD risk on auditory processing, criteria in defining the familial risk has varied (e,g., Leppänen et al., 2002; van Leeuwen et al., 2008). While criteria of 1.5 to 2 SD below population average in reading tests were often mentioned for DD (see diagnostic manual ICD-10, World Health Organization, 2016; Peterson and Pennington, 2015), parents in many previous infant MMR studies have only met a clearly looser criterion of, for example, 1 SD below average (Leppänen et al., 2002; van Leeuwen et al., 2008). Liberal criteria for DD risk increase the likelihood of null or random findings in group comparisons, as some or all participants in the DD risk group may actually not have a notable familial and genetic risk. To address this issue in the present study and to take into account the heterogeneity of the familial DD background in our DyslexiaBaby sample (Virtala et al., 2022), infants with different types of familial DD risk were treated as separate subgroups. Moreover, group comparisons were only conducted between the control group and the risk group demonstrating the clearest familial DD background (hereafter termed high risk group). Furthermore, in order to study the relevance of early phoneme discrimination skills for later language development irrespective of familial risk, we also studied the association of the phoneme-MMRs with subsequent language skills across the subgroups.

We hypothesized that phoneme changes in a sequence of acoustically varying speech sounds would elicit P-MMRs in newborn infants, in line with previous evidence (Dehaene-Lambertz and Pena, 2001). Based on leading theories on DD (Eden et al., 2016; Giraud and Ramus, 2013) and our previous adult findings (Virtala et al., 2020), we expected that infants in the high risk group face challenges in phonetic learning and would therefore show diminished phoneme-P-MMRs. Previous infant-MMR studies demonstrated atypical MMR lateralization in DD risk, or diminished amplitudes particularly at the left hemisphere (Leppänen et al., 2002; van Leeuwen et al., 2008; see also results in older children in Maurer et al., 2003). We therefore expected to see atypically bi- or right-lateralized phoneme-P-MMRs in the high risk group relative to those in the control group. Based on the aforementioned DD theories as well as on previous findings of infant-MMR amplitudes being associated with subsequent language and literacy (Cantiani et al., 2016; Oh et al., 2019; van Zuijen et al., 2013; Volkmer and Schulte-Koerne, 2018), we expected the MMRs to correlate with language skills at 28 months.

In addition to probing phoneme discrimination, the current paradigm allows for recording MMRs elicited by sound-order rule violations, reflecting implicit auditory rule extraction (for results in DD adults, see Virtala et al., 2021; for a comparable non-speech paradigm in neonates, see Carral et al., 2005). We briefly report these MMRs as well, however, these results should be considered as very preliminary, since only 8 infants in the control (non-risk) group (N=17 in the high DD risk group) had a sufficient number of trials for these violations. MMRs to rule violations, which are unfamiliar for the infants, should indicate implicit learning of the auditory rule (i.e., detecting the rule without explicit knowledge or instructions). We expected to find MMRs to the violations in the non-risk infants, in line with Carral et al. (2005), but no MMRs in the high-risk group based on the implicit or procedural learning deficit theory on DD (Krishnan et al., 2016; Ullman et al., 2020) and our recent adult findings (Virtala et al., 2021).

## 2. Methods

### 2.1 Participants

The total sample (a subsample of the DyslexiaBaby study; see recruitment and criteria in Thiede et al., 2019) consisted of 59 healthy newborn infants born full term (37–41 gestational weeks, birth weight > 2500 g) with normal hearing (confirmed with the optoacoustic emission test at the hospital after birth or, in three cases, by no indication of hearing problems during a three-year follow-up). One additional infant participated in the study but his/her EEG data were excluded due to poor quality (insufficient number of artefact-free trials, <30, for both deviant types). The study was approved by the local ethical review board and was conducted in compliance with the Declaration of Helsinki. Written informed consent was obtained from one or both parents of the infants at study enrollment.

Forty-three infants had a family history of DD (risk group) and 16 belonged to the control no-risk group (Table 1; language assessment scores missing from three control group infants). Due to great heterogeneity in the familial history of DD in the risk group, it was further divided into three subgroups. Those considered at *mild* (N=15) or *high* risk (N=18) had at least one biological parent with DD, confirmed by a recent diagnostic statement from a health care professional, or if it was missing, by researchers of the DyslexiaBaby study in a clinical interview and a standardized Finnish reading and writing skills test (Nevala et al., 2006). Parents in the mild risk subgroup were categorized by speed or accuracy one standard deviation below the norms, and parents in the high risk subgroup by two standard deviations below the norms in at least two subtests of the reading and writing skill test. Those considered at *low* risk (N=10) had at least one biological parent who, despite reporting childhood difficulties in reading and writing, performed within norms in the test, indicating compensated DD. Additionally, in order to ensure familial DD risk, the parent had to have a first-degree relative with DD. An infant was included in the control group if the parents (in one case one parent, since the other parent was unavailable) had not earlier been diagnosed with or been suspected of any language, reading, or learning disorders. Group comparisons for the MMR amplitudes to the phoneme changes were conducted to the two extreme groups, i.e., the control (N=16) vs. high risk group (N=18). Associations of the MMR amplitudes with language skills were studied across the whole sample (N=56; N=3 remaining participants not included due to missing language assessment data). For details, see 2.5 and 2.6.

**Table 1.** Demographic information of the groups in the final sample (standard deviation in parentheses). Apgar score (range 1–10) demonstrates the highest score of the infant 5/10 min. after birth. Parents in the low socio-economic status (SES) group have no higher (BA or higher) education.

|  | control N=16 |  | risk N=43 |  |
| --- | --- | --- | --- | --- |
|  |  | low risk<br>N=10 | mild risk<br>N=15 | high risk<br>N=18 |
| sex (f/m) | 8/8 | 7/3 | 8/7 | 9/9 |
| age, d | 11.5 (3.0) | 10.2 (3.3) | 9.7 (3.2) | 9.0 (3.4) |
| GA, w | 40.2 (1.4) | 40.0 (1.0) | 40.1 (1.4) | 40.1 (1.0) |
| Apgar | 9.5 (0.5) | 9.4 (0.5) | 9.5 (0.6) | 9.5 (0.6) |
| birth weight, g | 3643.4 (500.6) | 3579.7 (434.3) | 3680.6 (582.8) | 3563.9 (328.7) |
| birth height, cm | 51.0 (1.8) | 50.5 (1.5) | 50.8 (2.1) | 50.7 (1.6) |
| SES (low/high) | 2/14 | 3/7 | 2/13 | 1/17 |

*Note 1.* For three control group infants, language assessment data at 28 months was not available. Apgar score was missing from one infant in the high risk subgroup, but no complications during delivery were reported and the infant was considered to be healthy.
*Note 2.* Group comparisons for all subgroups and for the control vs. high risk group were conducted with Chi-Square and One-way Analysis of Variance (ANOVA) tests on the variables in the table. These tests yielded $p = .163$ for age between all subgroups and $p = .032$ for age between the control and high risk groups, and $p > .20$ for all other comparisons. Age did not correlate statistically significantly with the MMR amplitudes to vowel deviants, Pearson correlation $r(57) = .115$ , $p = .386$ , across the groups. For MMR quantification, see section 2.5.
*Note 3.* N=13 infants in the high risk group, N=10 infants in the mild risk group, and N=3 infants in the low risk group participated in a music listening intervention after the EEG recording, between birth and six months of age.

We also report preliminary results from a smaller sample of infants (control group N=8, low risk group N=9, mild risk group N=14, and high risk group N=17, Table 2) for the rule violation responses. The rule violation data of the rest of the infants (N=11, 8 from the control group and 3 from the risk groups) were excluded due to an insufficient number of artefact-free trials (<30), resulting from the lower total amount of presented rule violation trials compared to vowel deviant trials in the experimental paradigm.

**Table 2.**
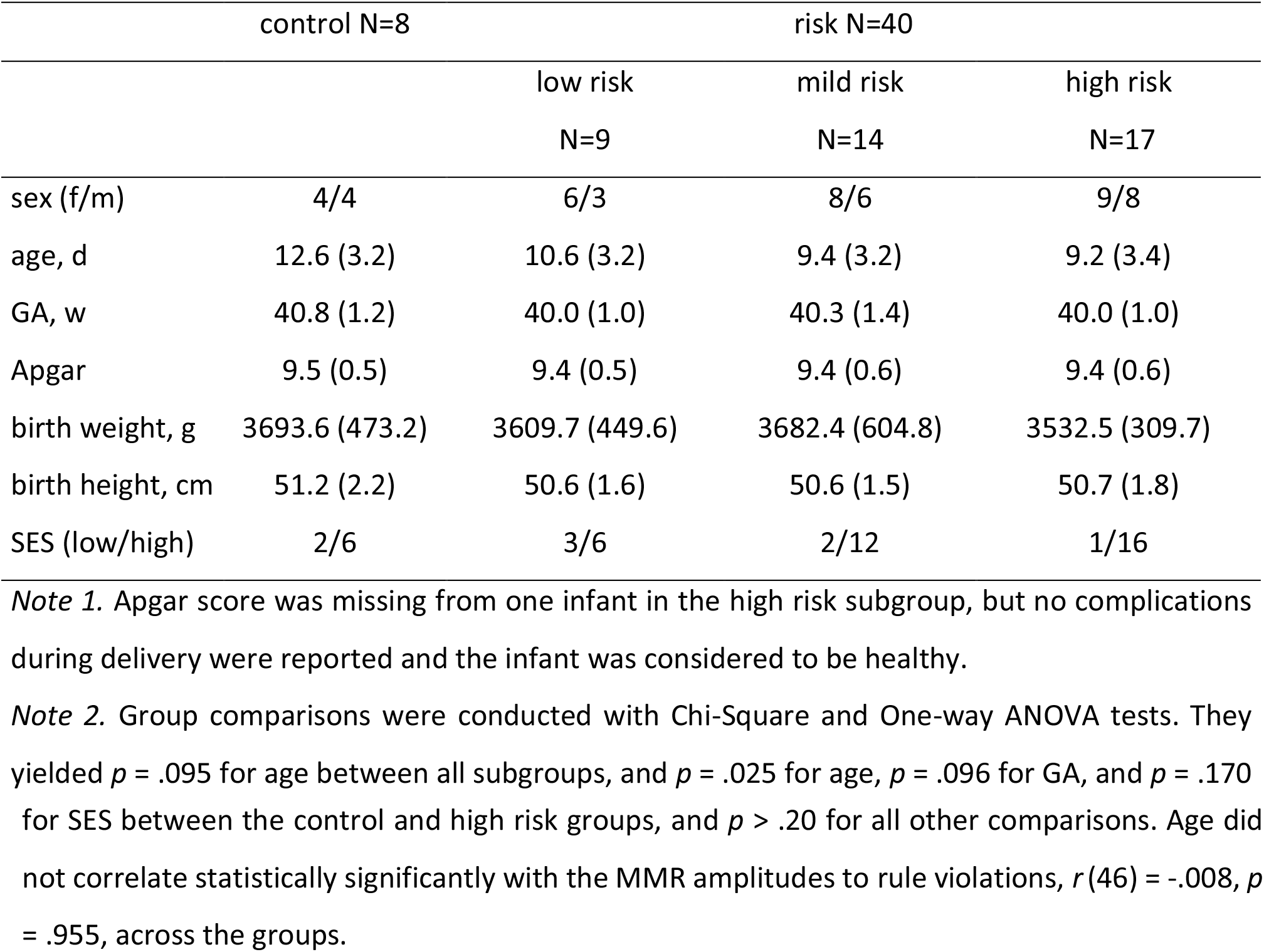
Demographic information of the groups in the rule violation data (standard deviation in parentheses). Apgar score (range 1–10) demonstrates the highest score of the infant 5/10 min after birth. Parents in the low socio-economic status (SES) group have no higher (BA level or higher) education.

It is noteworthy that N=26/43 of the risk group infants participated in a music listening intervention between birth and six months of age (for description, see Virtala and Partanen, 2018). Therefore, the language assessment scores at 28 months are not compared between groups in the present study: they are only analyzed in relation to the neonatal MMRs to determine whether these responses are associated with later language measures.

### 2.2 EEG stimuli and paradigm

The stimuli were pairs of Finnish phonemes /i/ and /æ/ uttered by a female native Finnish speaker, and edited using Praat 5.4.01 (Boersma and Weenink, 2014) and Adobe Audition CS6 5.0. Build 708 (Adobe Systems Inc., California, USA; stimuli and paradigm are described in Virtala et al., 2018). Sound intensity level between the phonemes was root-mean-square normalized. Phonemes (natural ending cut to a duration of 230 ms, fade-out between 190 and 230 ms) were transposed to seven *f_o_* levels (−4, −3, −2, −1, 1, 2, and 3 semitones from the original *f_o_* of ~205 Hz) and combined to 530-ms pairs, with a 70-ms silence in-between, with all possible /i/-/i/ and /i/-/æ/ combinations except that 1) there had to be at least two semitones between phonemes (thus rising-*f_o_* pairs could not start at the two highest, or descending-*f_o_* pairs at the two lowest *f_o_* levels), and 2) /æ/ always had a higher frequency than /i/ in /i/-/æ/ pairs (thus /æ/ with the two lowest *f_o_* levels and /i/ with the two highest *f_o_* levels not appearing in the stimuli).

In the 21-minute-long oddball sequence, /i/-/i/ pairs with a rising pitch served as the repeating standard (p = .8, 1008 stimuli) while /i/-/æ/ pairs (vowel deviants) and /i/-/i/ pairs with a falling pitch (rule violations) appeared as infrequent deviants (p = .1, 126 stimuli each). The pairs were presented in a pseudo-random order with at least one standard preceding a deviant. Different standard and deviant pairs appeared with equal probabilities. The pair onset-to-onset interval was 1000 ms with a 25-ms-jitter in 10 ms steps (975, 985… and 1025 ms) to reduce phase-locked neural activity.

### 2.3 EEG data acquisition

EEG was recorded at Jorvi Hospital of the Helsinki University Hospital in Espoo (n=51) and at the infant laboratory of the University of Jyväskylä, Finland (n=8, see supplemental material) with identical equipment (ActiCap headcap, QuickAmp 10.08.14 EEG amplifier, and BrainVision Recorder 1.20.0801; BrainProducts GmbH, Germany) and protocol. EEG (sampled at 500 Hz; 0-100-Hz bandpass filter) was recorded with 18 active electrodes placed according to the international 10/20 system (Fp1, Fp2, F7, F8, F3, F4, Fz, C3, C4, Cz, P7, P8, P3, P4, Pz, Oz, LM, RM) and online-referenced to the average of all electrodes.

A trained nurse or research assistant performed the recording (lasting 1–1.5 hours including the cap placement and removal) in a quiet hospital room (at Jorvi Hospital) or a sound-proof laboratory (at the University of Jyväskylä) and determined the state of the infant during the recording as ‘active sleep’, ‘quiet sleep’, ‘awake’, or ‘intermediate sleep stage’ (based on Grigg-Damberger et al., 2007). The infants lied on their back in a crib while the stimuli were presented (Presentation 17.2, Neurobehavioral Systems Ltd., USA) through a loudspeaker (Genelec speaker) at 40 cm from their head with an intensity of ≈65 dB SPL at the infant’s head. Data from all different sleep/alertness states were pooled for the analyses (for a similar protocol, see Thiede et al., 2019). All infants were reported to be in active sleep at some point during the experiment, while 16.7–33.3 % of the infants in each group were reported to be awake and 33.3–46.7 % of the infants in each group were reported to be in quiet sleep at some point during the experiment. There were no statistically significant group differences in these amounts in Chi-Square tests between all groups (proportion awake, *p* = .797; proportion quiet sleep, *p* = .873), or between the control and high risk group (proportion awake, p = .805; proportion quiet sleep, *p* = .435). The experiment reported here was carried out following another experiment (reported in Thiede et al., 2019), but only if the infant remained calm.

### 2.4 EEG data analysis

EEG was pre-processed with Matlab Releases 2017a–2020a (The MathWorks, Inc., USA) with the Toolboxes EEGLAB 14.0.0b and 2019_0 (Delorme and Makeig, 2004) and ERPLAB 7.0.0 (Lopez-Calderon and Luck, 2014). The continuous signals were prefiltered (0.025–40 Hz bandpass) and then visually inspected for bad electrodes. A maximum of five electrodes (28 %) were marked “bad” if they had flat or continuously noisy data. Peripheral (Fp1, Fp2, F7, F8, and Oz) bad electrodes were excluded from further analysis. Non-peripheral bad electrodes (F3, Fz, F4, C3, Cz, C4, P3, Pz, P4) were interpolated (max 2 per/infant, on average 1) using the rest of the valid electrodes. The data were first high-pass filtered (cut-off frequency 0.5 Hz, filter order 3300) and then low-pass filtered (cut-off frequency 25 Hz, filter order 264) using EEGLAB’s “eegfiltnew” function and FIR filter. The signals were then re-referenced to an average of LM, RM, P7, and P8 electrodes. An average of four electrodes close to each other was chosen as the reference in order to improve signal-to-noise ratio in the often poor-quality data in the peripheral electrodes, and to allow for referencing in situations where one of the reference electrodes was considered broken (in which case it and its contralateral pair were excluded from referencing) (see also Thiede et al., 2019, Kailaheimo-Lönnqvist et al., 2020).

The data were segmented into epochs of −100–840 ms around stimulus onset and baseline corrected by the average voltage in the −100–0 ms interval. Standard stimuli immediately following a deviant were omitted from the analyses. Epochs with their absolute amplitude exceeding ±120 μV at Fp1 and Fp2 were rejected to exclude eye-movement artifacts. For all electrodes included, epochs with a drift of > 100 μV or with data points exceeding ±3 SD from the mean response of all epochs were excluded (EEGlab’s jointprob-function, separately for each electrode and the response averaged across all electrodes). The remaining epochs were separately averaged for standard, vowel deviant, and rule violation stimuli. For the rule-violation ERPs, only the epochs with the phoneme pair starting at the middle four *f_o_* levels were included in the average (both standard and rule violation), resulting in a maximum trial amount of 60. This was done to equalize the *f_o_* levels appearing in the first phoneme of the standards and that of the rule violations, as the standards could not start from the two highest or rule violations from the two lowest *f_o_* levels (Section 2.2).

The final sample had on average 75 (range 33–106) accepted trials/infant for the vowel deviant, 456 (227–614) for the vowel standard, 38 (30–47) for the rule violation, and 215 (98–295) for the rule violation standard. Difference waveforms were separately calculated for the vowel deviants and rule violations by subtracting from the deviant response the corresponding standard response. For the difference waveforms, the baseline correction was shifted to −100–0 ms from the onset of the deviation, i.e., 200–300 ms from the pair onset.

### 2.5 ERP quantification and statistical analysis

To assess the MMR elicitation in each subgroup and to find the optimal measuring periods for the amplitude comparisons, standard and deviant ERPs were compared with cluster-based mass permutation tests (Fieldtrip toolbox with Matlab, Maris and Oostenveld, 2007; Oostenveld et al., 2011) from deviance onset (300 ms) to the end of the epoch (840 ms) at six fronto-central electrodes (F3, Fz, F4, C3, Cz, C4). We determined the latency windows showing significant differences (p < .05) between deviant and standard ERPs with the same polarity in adjacent time points at two or more neighboring electrodes. For each such significant cluster, the sum of *t*-values was computed, and the test statistic was defined as the maximum of the sum *t*-values. A null distribution for the test statistic was computed by permuting the deviant vs. standard stimulus labels 5000 times and calculating the test statistics for each iteration. The cluster sum *t*-values were deemed significant if they exceeded the top or bottom 2.5 percentile of the test statistics of the 5000 permuted iterations.

Based on these tests, and due to the broad and two-peaked response in the difference waveform (Figure 2), two amplitude measurement windows, consisting of significant clusters in at least some of the subgroups, 260–360 ms from deviance onset (early vowel-MMR) and 360–540 ms from deviance onset (late vowel-MMR) were chosen for the vowel-MMR in all subgroups. Mean amplitudes were calculated separately from the left (left region-of-interest, ROI: F3, C3) and right (right ROI: F4, C4) hemisphere electrodes. For the rule violation MMR, statistically significant clusters were found in the control group only. Its mean amplitudes were not calculated, as group comparisons were not conducted due to the small number of participants (N=8) in the control group.

**Figure 1.**
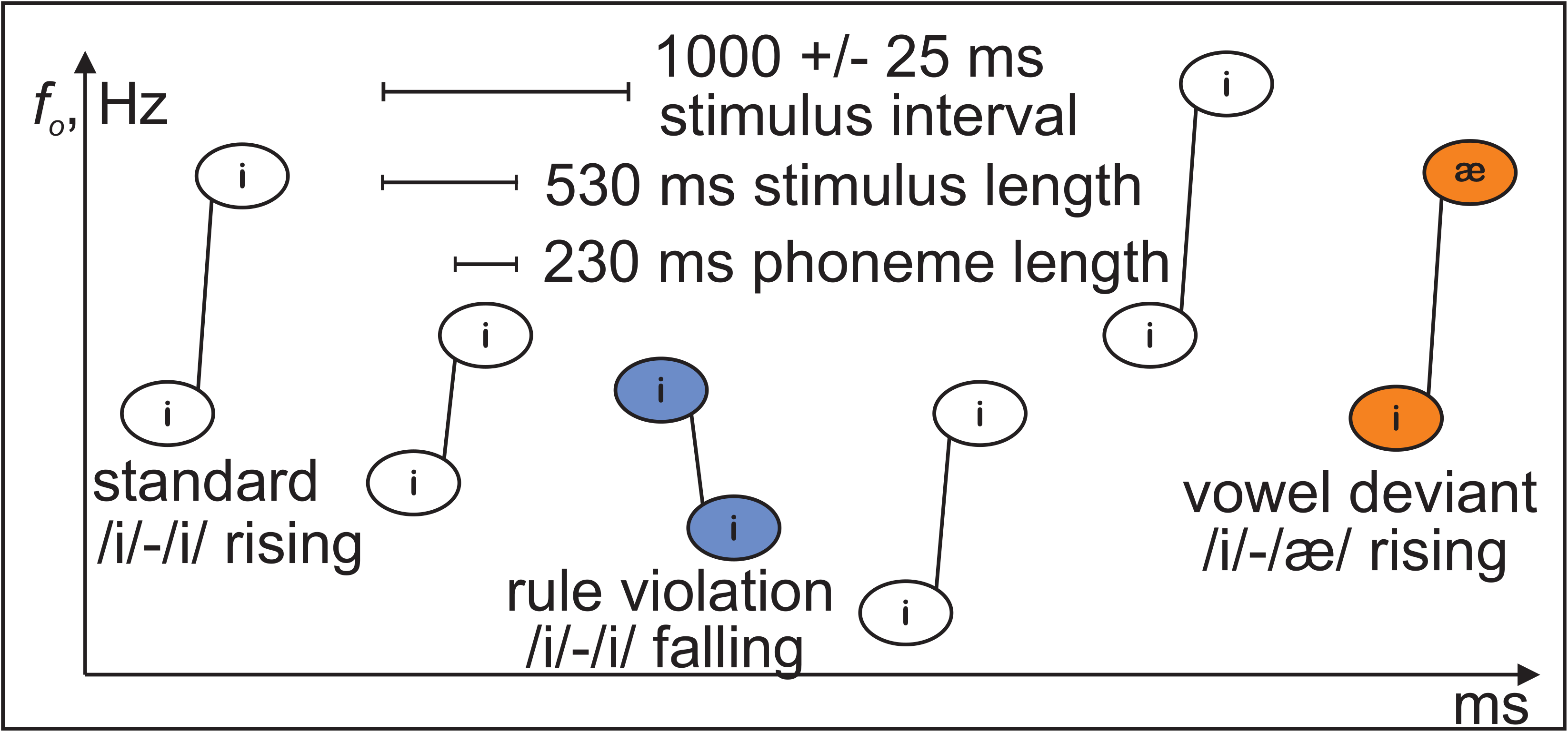
The experimental paradigm (adapted from Virtala et al., 2020).

**Figure 2.**
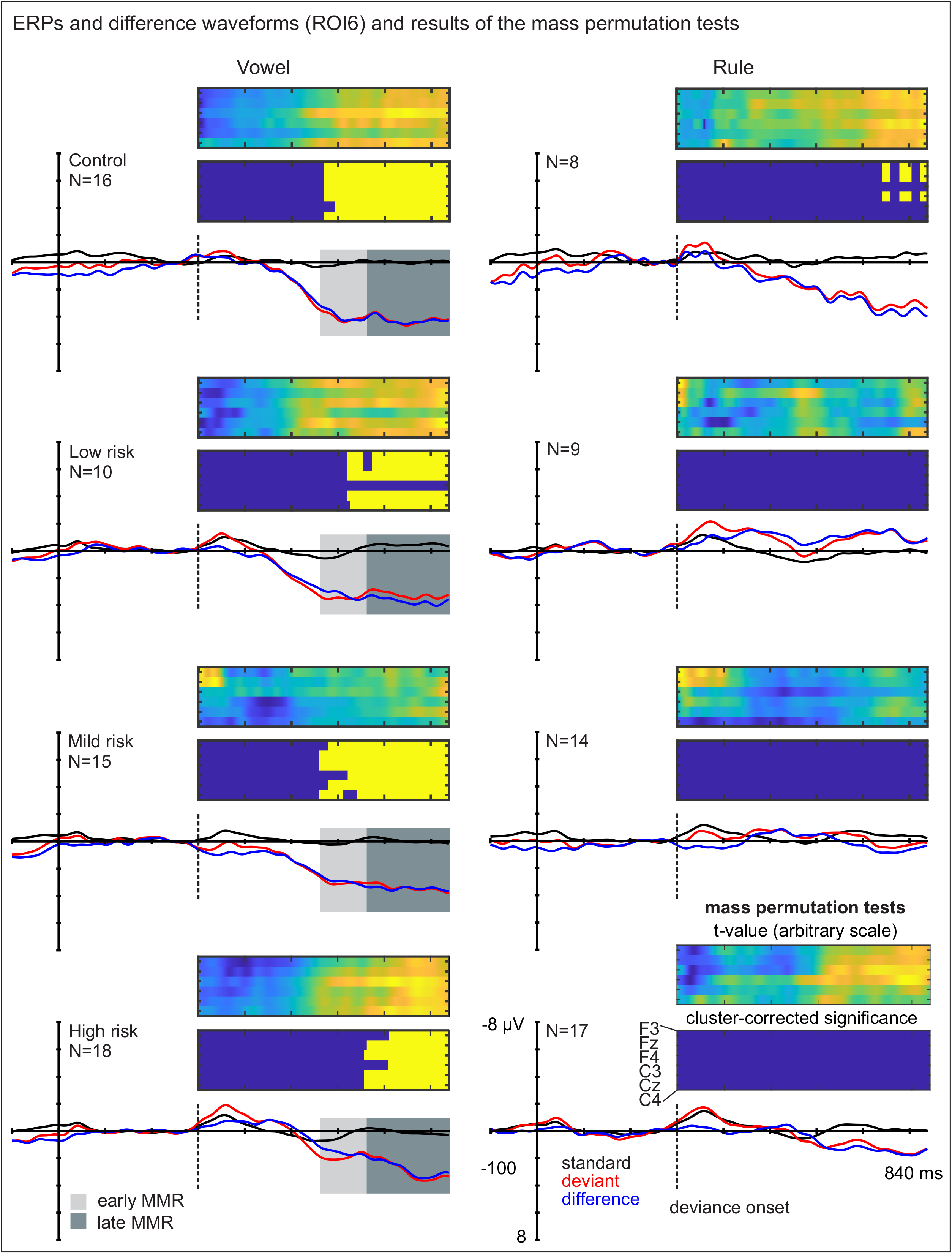
Standard (black) and deviant (red) ERPs and their difference (blue) at ROI6 in the four groups for the vowel deviant (left panel) and rule violation (right panel). The grey bars (light grey = early MMR, 260–360 ms from deviance onset; dark grey = late MMR, 360–540 ms from deviance onset) demonstrate the time windows used for vowel-MMR mean amplitude calculation from the difference waveform (light grey = early MMR, 260–360 ms from deviance onset; dark grey = late MMR, 360–540 ms from deviance onset). Mass permutation tests demonstrating the sum t values (top, blue indicating negative and yellow-orange positive difference) and significant clusters (bottom, corrected for multiple comparisons) of the standard vs. deviant ERPs at the six fronto-central electrodes. Yellow clusters mark statistically significant differences.

Due to the great heterogeneity in the degree and certainty of familial risk in the risk groups, only the control and high risk groups were included in the group comparison (for a similar protocol, see Thiede et al., 2019). Group differences and hemispheric lateralization of the vowel-MMR amplitude as well as their interactions were investigated using a Repeated-Measures ANOVA (RM-ANOVA) with hemisphere (left ROI vs. right ROI) and latency (early vs. late vowel-MMR) as within-subject’s factors and group (control vs. high risk) as a between-subject’s factor. Effect sizes are reported as partial eta squared (η^2^*_p_*), and Greenhouse-Geisser correction was applied when the assumption of sphericity was violated.

### 2.6 Language skill assessment at 28 months and statistical analysis

Language skill assessment at 28 months was conducted to N=56 children in the present sample (for N=3 in the control group, assessment was not completed due to scheduling issues or refusal to participate). The assessment took place in a behavioral laboratory space and was performed by a licensed speech and language pathologist or a master student of psychology or speech and language pathology under professional supervision. The researcher conducting the assessment was in most cases (N=49/56) unaware of the child’s group status (control group, risk group, or attendance to the music intervention). The assessment took approximately 2 hours. The parent was instructed to wait outside the laboratory, but if present in the room, they were advised not to help the child with the tests. Afterwards the child was rewarded with a small toy, and the parents were given oral feedback on the results.

The assessment consisted of two parts in counterbalanced order, 1) the complete Reynell Developmental Language Scale III (Reynell and Huntley, 2001) and 2) shorter test batteries or subtests as follows: Block Design and Object Assembly from Wechsler Preschool and Primary Scale of Intelligence III (Wechsler, 2003), Phonological Processing from NEPSY-II (Korkman et al., 2008), Rapid Automatized Naming (RAN; Puolakanaho et al., 2011), The Finnish Phonology Test (Kunnari et al., 2012), and a customized verbal short-term memory test. Only the Reynell test was included in the present analyses, as it was available from all children with very little disruptions during assessment reported for it compared to the other sub-tests. Moreover, the Reynell test is a commonly used and a well-established method for examining language skills in young children. It consists of two main scales, the Comprehension scale (CS) and the Expressive scale (ES), both with a number of subsections (10 for CS and 6 for ES) of increasing difficulty.

Associations between the raw scores of Reynell CS and ES and the early and late vowel-MMRs in the left and right ROIs were investigated with Spearman correlation analyses (due to the visually not normal distribution of the test scores) across the subgroups. The early and late MMRs were averaged for these analyses since they demonstrated similar group and hemispheric effects and to avoid the problems related to multiple testing. The whole sample was included to have a large N since behavioural test results at the age of 28 months are very unreliable and since in correlational analyses a high degree of variation is to be preferred. Both false discovery rate (FDR) corrected and not corrected results are reported.

## 3. Results

Vowel changes elicited statistically significant MMRs in all four subgroups (Figure 2, Table 3). Based on visual inspection of the waveforms, the vowel-MMRs were two-peaked, and in the permutation tests, the early vowel-MMR demonstrated significant clusters mainly in the control and mild risk groups, while the late vowel-MMR was statistically significant in all subgroups (Figure 2). Rule-MMR was statistically significantly elicited only in the control group, where significant clusters emerged in the left and middle electrode sites F3, Fz, and C3 towards the end of the epoch (Figure 2).

**Table 3.** Mean amplitudes (in μV) and standard deviations (SD) of the mismatch responses (MMRs) to the vowel deviants at the early and late latency on the left and right region-of-interest (ROI) in the four groups. The groups included in the RM-ANOVA are bolded.

| Latency | ROI | **Control** | Low risk | Mild risk | **High risk** |
| --- | --- | --- | --- | --- | --- |
| Early | Left | **3.58 (4.03)** | 3.33 (3.50) | 2.74 (3.56) | **1.15 (2.47)** |
|  | Right | **4.62 (3.78)** | 2.95 (3.01) | 3.21 (4.11) | **1.92 (2.66)** |
| Late | Left | **3.73 (3.47)** | 3.43 (3.77) | 3.23 (3.57) | **2.38 (2.88)** |
|  | Right | **5.13 (3.64)** | 3.50 (3.06) | 3.50 (4.12) | **3.16 (2.88)** |

The group comparisons, conducted on the high and no risk groups only, revealed statistically significantly larger vowel MMR amplitudes in the control (4.23 μV) than high risk group (2.15 μV), F(1,32)=4.299, *p*=.046, η^2^_p_ =.118 across the hemispheres and the early and late latencies, and in the right (3.71 μV) than left hemisphere (2.71 μV), F(1,32)=8.836, p=.006, η^2^_p_ =.216, across the two groups and the early and late latencies (Figures 3 and 4).

**Figure 3.**
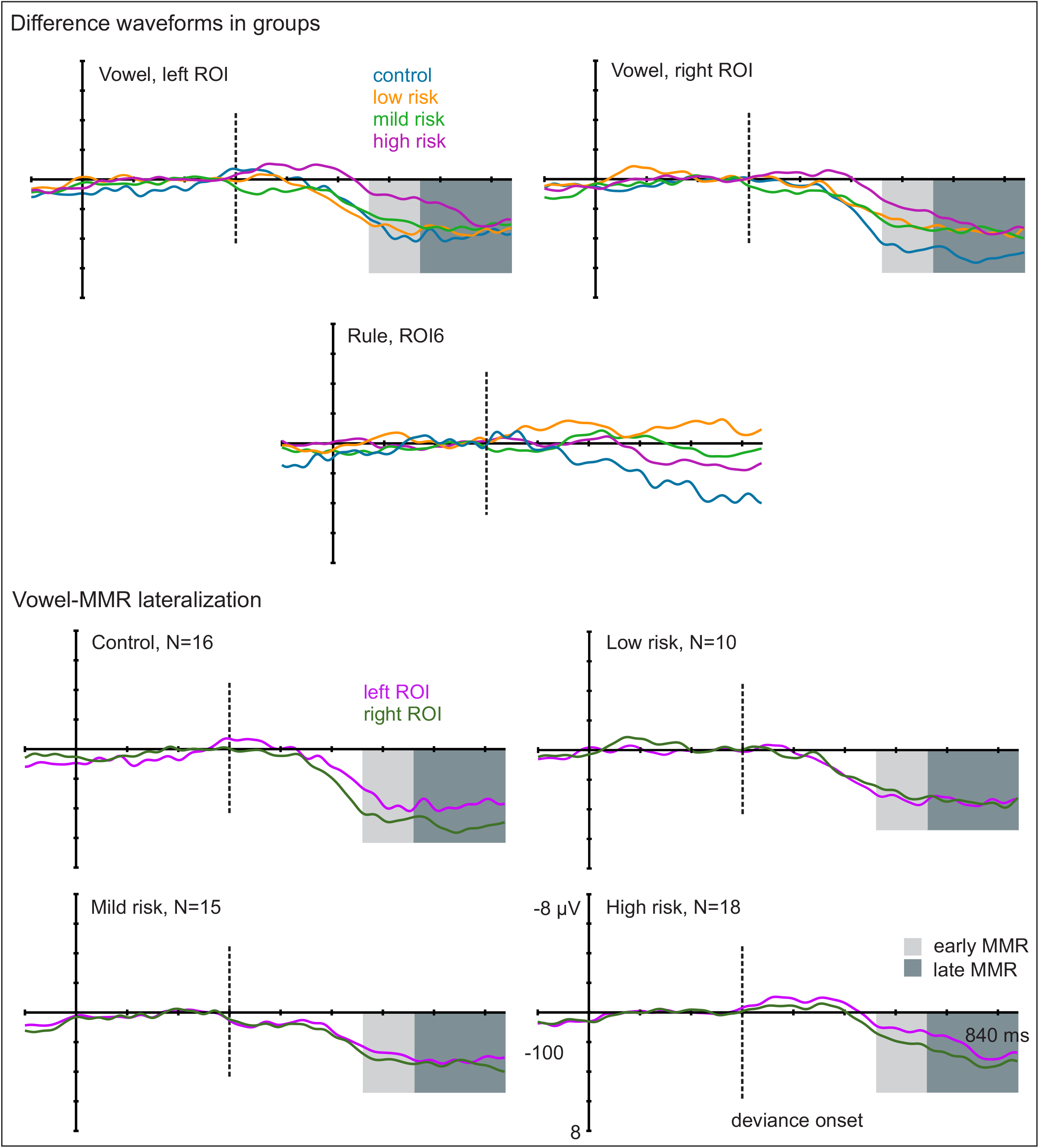
Top: Difference waveforms demonstrating the vowel-MMRs at left and right ROI and rule-MMRs at ROI6 in the four groups. Bottom: Vowel-MMR difference waveforms in the left and right ROIs in the four groups, with the grey bars demonstrating the time windows used for mean amplitude calculation (light grey = early MMR, 260–360 ms from deviance onset; dark grey = late MMR, 360–540 ms from deviance onset).

**Figure 4.**
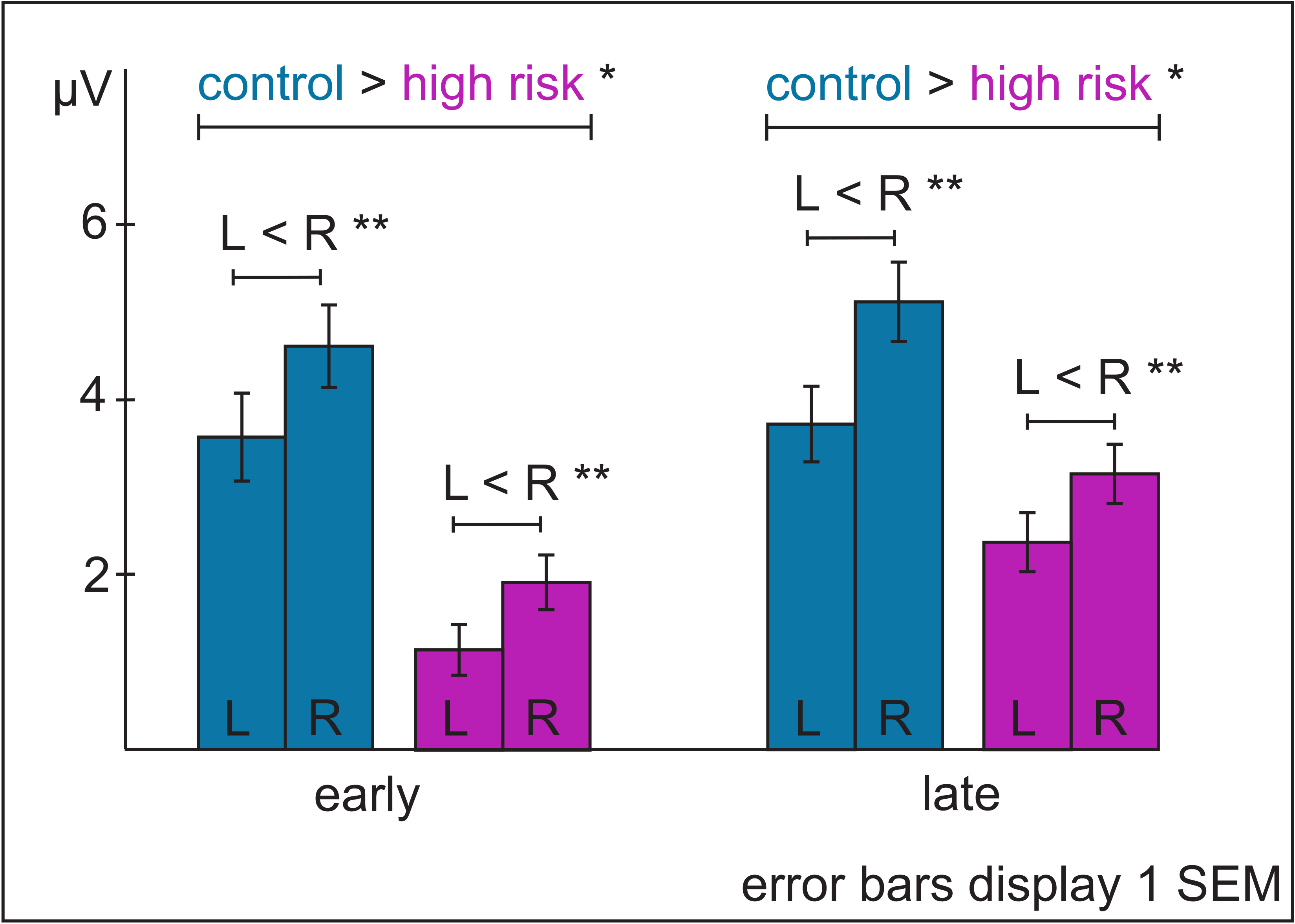
The mean amplitudes and their standard errors of mean (SEM) of the early and late vowel-MMRs on the left (L) and right (R) region-of-interest in the control and high risk group. Results of the RM-ANOVA are marked with asterisks (* p < .05, ** p < .01).

The correlation analyses were conducted over the whole sample. As the early and late vowel-MMRs demonstrated similar group and hemispheric effects and in order to reduce the problems of multiple testing, an average of the early and late vowel-MMRs was calculated separately for the right and left ROIs. Across the subgroups, vowel-MMR in the right ROI demonstrated a negative correlation with Reynell ES, *r*(54)=-0.323, *p*=.015, but the correlation did not remain statistically significant after correcting for multiple comparisons (FDR-corrected critical value, p=0.0125).

We also calculated a laterality index (LI) for the vowel-MMR again averaged across the early and late latencies: (left ROI – right ROI) / (left ROI + right ROI); for a similar protocol, see Seghier (2008). This index gives a more straight-forward measure of the ratio of activations measured on the left and right scalp, helping to interpret the relative contributions of these hemispheres in stimulus processing. The LI gives values between −1 and 1, with positive values when ROI left > ROI right and negative values when ROI right > ROI left, but only when amplitude values at both ROIs are positive. Negative amplitude values in individual participants were interpreted to indicate the absence of a P-MMR. Data of the participants with negative amplitudes on either of the two ROI’s were therefore excluded from the LI calculation (N=16, N=4 from the control group and N=12 from the risk groups), resulting in N=40 participants with both LI and language score data. The vowel-MMR LI correlated statistically significantly with Reynell ES, *r*(38)=.453, *p*=.003 (FDR-corrected critical value *p* =.025), indicating that a left-lateralized vowel-MMR was associated with higher expressive language scores (Figure 5).

**Figure 5.**
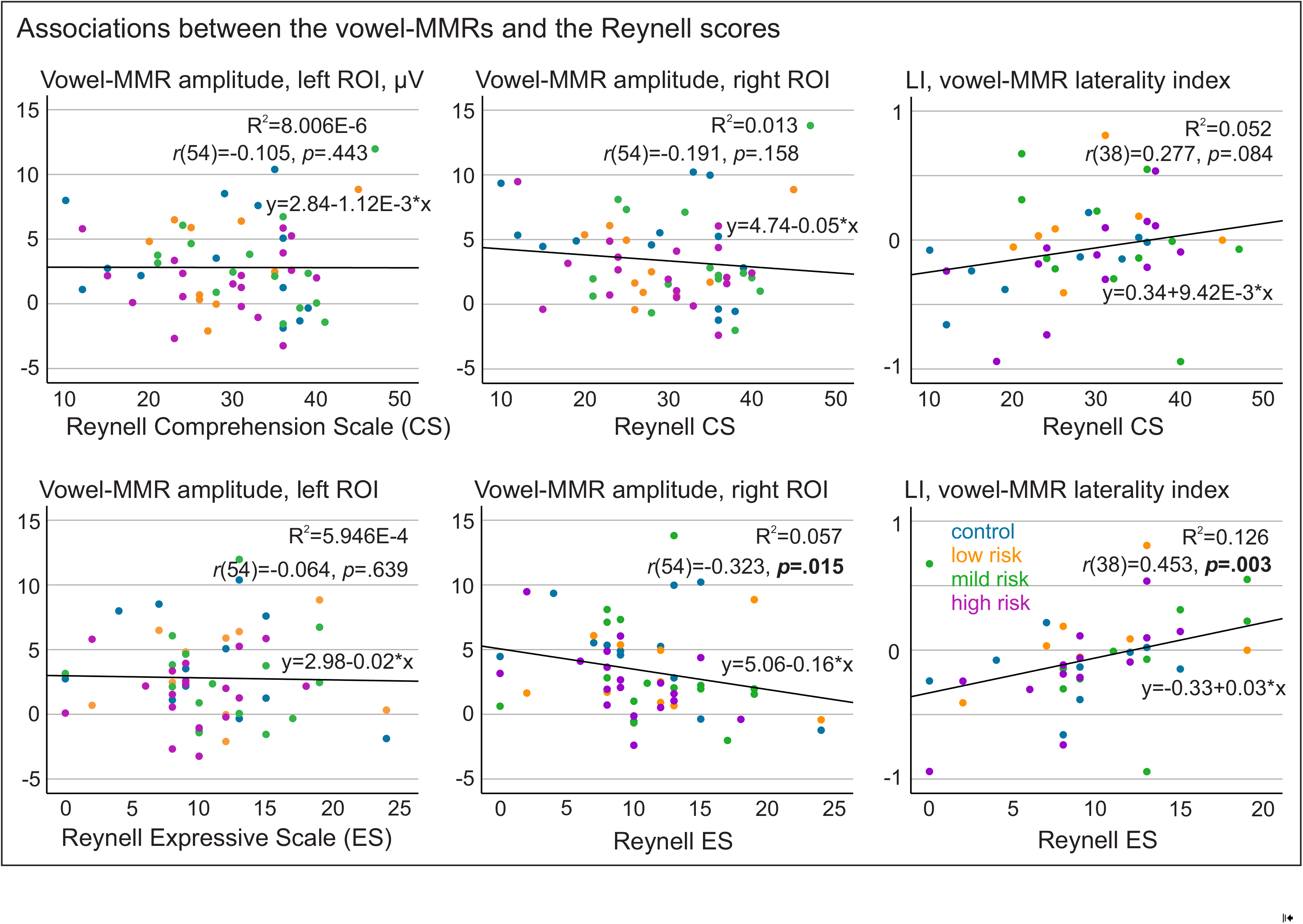
Scatterplots illustrating the Spearman correlations (groups pooled) of the vowel-MMR mean amplitudes in the left and right ROI and the vowel-MMR laterality index with raw scores of the Reynell Comprehension scale (CS, top panel) and Expressive Scale (ES, bottom panel). Dots where LI > 0 indicate left-lateralized MMRs. Bolded p-values indicate statistical significance (*p* < .05) before correcting for multiple comparisons; only the correlation between the LI and Reynell ES remained statistically significant after corrections.

## 4. Discussion

According to prevailing views, learning to perceive sound relationships underlying phoneme identities is essential for early phonetic learning and language acquisition (Kuhl, 2010), and a phonological deficit is central in the development of DD (Vellutino et al., 2004). Based on these theories, we examined phoneme discrimination abilities in newborns with or without a risk for DD utilizing auditory ERPs. We compared MMRs elicited by phoneme changes in an acoustically variable context between control and high DD risk groups. In addition, we studied the associations of the MMRs with subsequent language skills. As hypothesized, we found statistically significant MMRs to phoneme changes in all participant groups irrespective of DD risk, but diminished amplitudes in the infants who were at high DD risk due to moderate or severe parental DD symptoms. The phoneme-MMR had a right-hemispheric lateralization pattern across groups. Its left-hemispheric lateralization was associated with better expressive language scores. Moreover, we found preliminary results suggesting that sound-order rule violations embedded in the stimulus sequences elicited statistically significant MMRs only in the control group. As the control group had a very small sample size (N=8) for the rule-MMR data (due to which these data were not analyzed further), this finding should be treated with great caution. Together, the results demonstrate auditory abilities in newborns that are potentially relevant for language acquisition. The group differences in MMRs and their elicitation suggest that deficits in phoneme discrimination and, possibly, implicit auditory rule extraction may play a role in (or at least accompany) DD risk. The associations of MMRs to language skills highlighted the relevance of early phoneme extraction abilities, likely reflecting phonetic learning, for subsequent language development.

The phoneme-MMRs elicited robustly in all groups suggest that newborns can detect vowel changes even when the speech signal has *f_o_* variation, mimicking, e.g., prosodic variation in natural speech. The finding corroborates those of a previous study showing MMRs to stop consonant changes in syllables spoken by four different speakers (Dehaene-Lambertz and Pena, 2001). The results may reflect pre- or postnatal phonetic learning, or alternatively, they may indicate readiness to rapidly categorize the speech sounds to two phoneme classes during the experiment. Overall, these results imply that infants have a readiness to pre-attentively extract phonetically relevant information for acquiring phoneme categories already at birth.

The diminished MMR to phoneme changes in newborns with high DD risk compared to controls suggests an early deficit in accurately discriminating the phonemes, consistent with previous studies which, however, used repetitive, acoustically non-variable stimulus streams (van Leeuwen et al., 2008, 2006). Our stimuli with acoustic variation provide, relative to those studies, a speech sound context more closely resembling natural speaker variation. In our previous study on adults with these same stimuli, diminished MMN/N2b responses were found in DD, and the group difference vanished when stimuli were presented in a repetitive context (without *f_o_* variation; Virtala et al., 2020). Similarly, in our previous study with partly the same infants as in the present study, a P-MMR to vowel changes in a repetitive pseudo-word context did not show statistically significant differences between control and high risk groups (Thiede et al., 2019). Together these findings suggest that DD and its familial risk are not always associated with neural auditory deficits in simple, repetitive contexts that may rely on basic acoustic discrimination skills. Still, DD (risk) may compromise the ability to group auditory stimuli into categories based on their shared acoustic features, which is likely important for phonetic learning, that is, learning the native language phoneme categories.

However, in the absence of a continuum from one phoneme to another, the present paradigm should not be considered a measure of categorical phoneme processing. Nevertheless, our results are in line with behavioral evidence of deficient categorical phoneme processing associated with DD (Noordenbos and Serniclaes, 2015). A phoneme categorization deficit in infants could lead to language learning dysfunctions, since efficient acquisition of native language phonemes during infancy was proposed to be vital for good language development (Kuhl, 2010) and for eventually mapping the phonemes with their written input, i.e., literacy acquisition (Serniclaes, 2018). Indeed, as discussed below, phoneme-MMRs in the present study showed associations to subsequent language development. In the future, contrasting the results obtained with our ERP paradigm with a behavioral categorical phoneme discrimination or differentiation task would provide important information about the nature of the process tapped in the present study.

In the present study, somewhat unexpectedly, right-hemispheric lateralization of the phoneme-MMRs was seen in the newborns across groups. While most previous studies have reported left-lateralized phoneme discrimination (Kuuluvainen et al., 2016; Sorokin et al., 2010), this has not always been the case in newborn studies (Perani et al., 2011; Thiede et al., 2019), and it is possible that language processing becomes predominantly left-lateralized only later on (Olulade et al., 2020). Indeed, a recent study suggests that the right hemisphere is involved in language processing in young children, but that its contribution declines with age (Olulade et al., 2020; see also Dehaene-Lambertz et al., 2002).

We found a positive correlation between the left-lateralization of the phoneme-MMR and expressive language test scores at 28 months. These findings are in line with our hypotheses and previous evidence (Cantiani et al., 2016; van Zuijen et al., 2013; Volkmer and Schulte-Koerne, 2018). In both infants (Guttorm et al., 2010, 2005) and older children (Kuuluvainen et al., 2016; Maurer et al., 2009), left-lateralized or left-hemispheric neural responses have been shown to have positive associations to concurrent or subsequent language and literacy. Also, most previous studies have found positive associations of left-rather than right-hemispheric MMRs with language abilities (e.g., Leppänen et al., 2002).

Data of several infants (~1/4 of the sample) unfortunately had to be excluded from the laterality index analyses due to negative MMR values that suggested absent P-MMRs in these infants and would have compromised the interpretation of the laterality index. While newborn MMRs may have both positive and negative polarities (e.g., Thiede et al., 2019), the phoneme-MMRs were calculated from a time window that showed a positive MMR in all subgroups, and no sign of statistically significant negative MMRs were seen in the group-wise permutation analyses (Figure 2). Also in our previous infant-MMR study, vowel deviants elicited P-MMRs only (Thiede et al., 2019), and negative MMRs were often reported in previous work from an earlier latency than what was used in the present study (e.g. 50–250 ms from deviance onset in Fellman et al., 2004; ~100 ms from deviance onset in Thiede et al., 2019 vs. 260–540 ms from deviance onset in the present study). For these reasons, we chose to exclude negative values from these analyses instead of including them in the laterality index or analyzing them separately.

Our stimulus paradigm included also violations of a rule according to which the first sound in the vowel pairs had a lower *f_o_* than the second one. Due to a low number of participants with these data (N=8 for the control and N=17 for the high risk group), we only tested whether the MMR to this violation was significantly elicited in each group and conducted no further analyses. The preliminary results, which have to be dealt with very cautiously, showed statistically significant P-MMRs to the rule violations in the control group, which are consistent with previous results obtained with nonspeech stimuli (Carral et al., 2005; Háden et al., 2015; Stefanics et al., 2009). Elicitation of the rule-MMR in the control infants suggests that they were able to detect a second order sound relationship, i.e., infrequent changes in the direction of pitched contour despite the *f_o_* variation. As the rule was novel, and yet elicited an MMR in the mostly sleeping newborns, we propose that this might reflect implicit statistical auditory learning abilities, possibly important for subsequent language development. The non-significant MMR to the rule violation in the DD risk groups might suggest difficulties in implicit rule extraction, previously shown in adults with DD (Virtala et al., 2021; see also behavioral findings Gabay and Holt, 2015; Gabay et al., 2015). Such rule-based auditory processing is essential for language learning, as language comprises of rules and regularities that are implicitly adopted during early development through statistical learning (Kuhl, 2010). Our preliminary results might indicate that infants at DD risk have a deficiency in this vital, implicit language-learning ability. Although DD is also associated with a general pitch processing deficit, it is unlikely to explain the absent rule-MMR in the DD risk groups: the deficit is typically seen with smaller pitch differences (< 10%) than the ones presented within the sound pairs of the present study (12–47%; Hämäläinen et al., 2013). However, as already stated, these results are compromised by the low number of infants from whom rule-MMRs were obtained (especially in the control group). This also hindered further analysis of group differences and associations to language skills, and therefore no firm conclusions can be drawn on the possible effects. Therefore, these very preliminary results await confirmation from future studies with higher participant numbers.

An attempt was made in the present study to deal with the heterogeneity and often liberal criteria in the definition of DD and DD risk, by dividing the risk group to subsamples based on the expected degree of familial risk. Among the familial risk group, phoneme-MMR amplitudes of only those infants who had the clearest familial risk for DD, i.e., parental DD symptoms currently reaching the −2SD criterion used in ICD-10 (World Health Organization, 2016), were compared against the MMRs of the no-risk control group. To our knowledge, different DD (risk) criteria have not been compared in infant (or even adult) ERP studies. We hope that future studies employing large samples in longitudinal settings will pay increasing attention to these issues.

The current results provide evidence on the complex auditory abilities of newborn infants. Based on the observed associations to subsequent language scores and the diminished phoneme-MMRs in the high DD risk group, these abilities are likely to contribute to phonological development and language acquisition. As our results show, even complex auditory functions crucial for speech processing can be investigated already at birth. In the future, methodological advancements could enable the infant auditory ERPs to become more reliable and replicable also at the individual level. Identification of those aspects of auditory processing that serve as the best neural predictors of future language and reading problems could then help in targeting early support. If these early auditory deficits have a causal role in language and literacy development, interventions targeted to ameliorate them could also prevent later problems.

## Data availability statement

As no common repositories for neurocognitive data are available for the authors and the ethical permission does not include a clause determining the specifics of data availability, the data can be provided for readers upon reasonable request to the corresponding author, as is common practice in the field.

## Acknowledgements and funding statement

The authors thank all families for their participation, research nurses Tarja Ilkka and Svetlana Permi for conducting the majority of the EEG recordings, Dr. Vesa Putkinen for his valuable advice in statistical data analysis, and all research assistants for their involvement in the project. The authors have received funding from the Academy of Finland (grant numbers 276414, 316970, 346211), Jane and Aatos Erkko Foundation, Finland, Kela (The Social Insurance Institution in Finland), Finnish Cultural Foundation, and the National Research Fund of Hungary (grant number K115385).

## Ethics approval statement

These data were collected as part of the DyslexiaBaby longitudinal study, approved by the Ethics Committee for Gynaecology and Obstetrics, Pediatrics and Psychiatry of the Hospital District of Helsinki and Uusimaa and in compliance with the Declaration of Helsinki.

## Author statement

**Paula Virtala:** Participation in conceptualization, Methodology, Software, Formal analysis, Investigation, Writing – original draft, Writing – review & editing, Visualization, Supervision, Project administration. **Teija Kujala:** Conceptualization, Methodology, Resources, Writing – review & editing, Supervision, Project administration, Funding acquisition. **Eino Partanen:** Participation in conceptualization, Methodology, Software, Formal analysis, Investigation, Writing – review & editing. **Jarmo Hämäläinen:** Investigation, Writing – review & editing. **Istvan Winkler:** Writing – review & editing.

## Supplementary Material

Supplemental_material_Virtala_Kujala.docx

