## Supplemental_material for "Neural phoneme discrimination in variable speech in newborns – associations with dyslexia risk and later language skills"

*Comparison between recording sites*

In order to ensure that the research environment (Jorvi/Jyväskylä) did not influence the results obtained, the statistical analyses were repeated with the N=51 infants recorded in Jorvi (i.e., N=8 infants recorded in Jyväskylä were excluded, N=6 of these from the control group, N=1 from the low risk group, and N=1 from the mild risk group).

The RM-ANOVA comparing the control (N=10) and high risk group (N=18) still demonstrated statistically significantly larger vowel-MMR amplitudes on the right than left hemisphere, $F(1,26) = 10.001$, $p = .004$, but the main effect of group no longer reached significance ($p = .356$), although the control group still demonstrated numerically larger MMR amplitudes than the high risk group (3.17 *vs.* 2.15 µV, respectively).

The negative correlation between the vowel-MMR (an average of the early and late vowel-MMRs) at the right ROI with the Reynell ES across groups remained statistically significant, $r(46) = -.414$, $p = .003$, while a statistically nearly significant correlation to Reynell CS was also obtained, $r(46) = -.247$, $p = .091$ (Bonferroni-corrected criterion over four tests $p = .013$). The positive correlation between the vowel-MMR LI and Reynell ES also remained statistically significant, $r(31) = .545$, $p = .001$ (Bonferroni-corrected criterion over two tests $p = .025$), indicating an association between left-lateralized vowel-MMRs and larger Reynell ES scores. The correlation between the vowel-MMR at the left ROI with the Reynell ES, when the infants demonstrating negative amplitudes on either hemisphere were excluded, remained positive but no longer reached statistical significance, $r(31) = .299$, $p = .091$ (Bonferroni-corrected criterion over four tests $p = .013$).

To conclude, likely due to the small sample sizes and the high proportion of control group infants among those infants that were recorded in Jyväskylä (N=6/8), the group difference obtained between control and high risk groups in vowel-MMR amplitudes did not hold in the present data when the Jyväskylä infants were excluded. Further studies should confirm whether the group difference holds with other, larger samples. However, the results on correlations between MMRs, their lateralization, and the subsequent language skills remained robust also within the Jorvi recording site.
